# ABIOTIC STRESS GENE 1 mediates aroma volatiles accumulation by activating MdLOX1a in apple

**DOI:** 10.1101/2022.02.24.481825

**Authors:** Jing Zhang, Susu Zhang, Yongxu Wang, Shuhui Zhang, Wenjun Liu, Nan Wang, Hongcheng Fang, Zongying Zhang, Xuesen Chen

**Affiliations:** State Key Laboratory of Crop Biology, College of Horticulture Sciences and Engineering, Shandong Agricultural University, Tai’an, 271018, Shandong, China; Xinjiang Production and Construction Corps Key Laboratory of Special Fruits and Vegetables Cultivation Physiology and Germplasm Resources Utilization, Department of Horticulture, College of Agriculture, Shihezi University, Shihezi, 832003, Xinjiang, China; State Forestry and Grassland Administration Key Laboratory of Silviculture in the Downstream Areas of the Yellow River, College of Forestry, Shandong Agricultural University, Tai’an, 271018, Shandong, China

**Keywords:** Apple, Lipoxygenase, MdLOX1a, ABIOTIC STRESS GENE 1, Aroma, Ester, Volatiles, Salt stress

## Abstract

Fruit aroma is an important organoleptic quality, which influences consumer preference and market competitiveness. Aroma compound synthesis pathways in plants have been widely identified of which the lipoxygenase pathway is crucial for fatty acid catabolism to form esters in apple. However, the regulatory mechanism of this pathway remains elusive. In this study, linear regression analysis and transgene verification revealed that the lipoxygenase MdLOX1a participates in ester biosynthesis. Yeast one-hybrid library screening indicated that a novel abiotic stress gene, *MdASG1* (*ABIOTIC STRESS GENE 1*), was a positive regulator of the *MdLOX1a* promoter and ester production based on yeast one-hybrid and dual-luciferase assays, and correlation analysis among eight apple cultivars. Overexpression of *MdASG1* in apple and tomato stimulated the lipoxygenase pathway and increased the fatty acid-derived volatile content, whereas the latter was decreased by *MdASG1* silencing. Furthermore, *MdASG1* overexpression enhanced the salt-stress tolerance of tomato and apple ‘Orin’ calli accompanied by a higher content of fatty acid-derived volatiles compared with that of non-stressed transgenic tomato fruit. Collectively, these findings indicate that *MdASG1* activates *MdLOX1a* expression and participates in the lipoxygenase pathway, subsequently increasing the accumulation of aroma compounds especially under moderate salt stress treatment. The results also provide insight into the regulation of aroma production, and the potential strategy of prudent development and utilization of saline-alkali land to produce high-quality fruit, thereby reducing pressure on arable land and ensuring national food security.

**One-sentence Summary:** MdASG1 directly activates *MdLOX1a* expression to promote aroma volatiles accumulation especially under moderate salt stress.

## INTRODUCTION

Plant volatile organic compounds are secondary metabolites that play important roles in biotic and abiotic stress responses, and act as signals to attract or repel pests, confer resistance to pathogens, and participate in seed dispersal (Rodriguez et al., 2013). Many volatiles are produced by plants at different developmental stages, especially during fruit ripening. A large number of volatile esters are produced by strawberry (*Fragaria vesca*), banana (*Musa sapientum*), apple (*Malus domestica*), and peach (*Prunus persica*) (Beekwilder et al., 2004; Souleyre et al., 2014; Cao et al., 2021). Fruit quality mainly reflects fruit shape, size, color, aroma, acidity, sugar content, and nutritional content. Among these traits, aroma is an important determinant of the commercial value of fruit. However, breeders tend to focus on yield, disease resistance, and fruit color, and pay little attention to flavor and thereby weaken customer motivation to buy apple fruit (Klee and Tieman, 2018). Therefore, improvement in fruit flavor is desirable to meet consumer demand.

The aroma compound synthesis pathway has been extensively studied in plants. Fruit esters are produced mainly from the fatty acid pathway contributing to straight-chain ester synthesis and the amino acid pathway contributing to branched-chain ester formation. In tomato (*Solanum lycopersicum*), a number of fatty acid-derived chemicals, including C5 or C6 aldehydes and alcohols, are formed in the lipoxygenase pathway (Stone et al., 2010). In apple, the β-oxidase and lipoxygenase pathways are the two main enzyme systems involved in fatty acid catabolism to form esters (Rowan et al., 1999). In the lipoxygenase pathway, lipoxygenases (LOX) catalyze polyunsaturated fatty acids, including linolenic and linoleic acid, to produce hydroperoxides (Feussner and Wasternack, 2002). The hydroperoxides are then converted to short-chain aldehydes and an oxo-acid by hydroperoxide lyase, which belongs to the cytochrome P450 superfamily (Matsui, 1998; Schwab et al., 2008). The short-chain aldehydes are further reduced to corresponding alcohols by alcohol dehydrogenase during fruit ripening (Manriquez et al., 2006; Schwab et al., 2008). Lastly, alcohol acyl-transferases (AATs) catalyze the acid donor, acyl-coenzyme A (acy-CoA), and alcohol acceptor to synthesize esters (Dunemann et al., 2012). Lipoxygenases are a non-heme iron-containing dioxygenase, which are classified as either 9-LOX or 13-LOX according to the position of the carbon targeted for oxygenation in the polyunsaturated fatty acid (Feussner et al., 2001). Lipoxygenases are also classified as type 1 or type 2 LOXs according to the sequence similarity. Lipoxygenases of tomato, pepino (*Solanum muricatum*), and kiwifruit (*Actinidia deliciosa*) are involved in aroma compound synthesis (Chen et al., 2004; Zhang et al., 2006; Zhang et al., 2009; Contreras et al., 2017).

Certain other factors affect the accumulation of fruit flavor compounds, including genetic differences (Kakiuchi et al., 2007), crop management (Mpelasoka and Behboudian, 2002), harvest date (Song and Bangerth, 1996), storage environment (Harb et al., 2012), and the plant hormones ethylene, abscisic acid (ABA), and jasmonic acid (JA) (Yang et al., 2016; Wu et al., 2018; Luo et al., 2021). Recently, transcriptional regulation of specific genes involved in aroma synthesis has been investigated. The ETHYLENE-INSENSITIVE3-LIKE (EIL) and NAC transcription factors activate terpene synthase gene *AaTPS1* transcription to control monoterpene production in kiwifruit (*Actinidia arguta*) (Nieuwenhuizen et al., 2015). NAC transcription factors modulate ester biosynthesis by regulating expression of the structural genes *FAD1* and *AAT10* in kiwifruit (Zhang et al., 2020; Wang et al., 2022) and activate *AAT* expression in tomato, peach, and apple (Cao et al., 2021). The AP2/ERF transcription factors EREB58, CitAP2.10, and CitERF71 may *trans*-activate the terpene synthase *TPS* to promote the synthesis of terpenes (Li et al., 2015; Shen et al., 2016; Li et al., 2017). Strawberry ethylene response factors FaERF9 and FaMYB98 form a protein complex, which indirectly activates strawberry quinone oxidoreductase (FaQR) expression, thereby promoting the synthesis of furanone (Zhang et al., 2018). The R2R3 MYB transcription factors FaEOBⅡ and FaDOF2 synergistically regulate the volatile phenylpropanoid pathway in strawberry (Medina-Puche et al., 2015; Molina-Hidalgo et al., 2017). In tomato, the MADS box transcription factor RIN and SlMYB75 directly bind to the promoter of genes for aroma compound synthesis pathway-related enzymes to activate their expression (Qin et al., 2012; Jian et al., 2019). In addition, other transcription factors, including a basic helix-loop-helix transcription factor (MYC2), *PRODUCTION OF ANTHOCYANIN PIGMENT 1* (*PAP1*), and basic leucine zipper (bZIP), mediate aroma compound biosynthesis (Hong et al., 2012; Zvi et al., 2012; Guo et al., 2018).

*ABIOTIC STRESS GENE 1* (*ASG1*) is an abiotic stress gene identified in *Solanum tuberosum* and *Arabidopsis thaliana* inducible by stress via an ABA-dependent pathway (Batelli et al., 2012). However, little information is available on whether *ASG1* mediates other biological activities, including aroma regulation. Stress can induce the production of secondary metabolites to improve fruit quality. Treatment with ABA reduces tannin content and positively affects grape (*Vitis vinifera*) fruit quality (Lacampagne et al., 2009). Abscisic acid drives the accumulation of secondary metabolites contributing to fruit aroma in grape and strawberry (Ferrandino and Lovisolo, 2014; Kadomura-Ishikawa et al., 2015). MdAREB2 is responsive to ABA and promotes soluble sugar accumulation by activating the expression of amylase and sugar transporter genes (Ma et al., 2017). Soil water stress can improve fruit quality by increasing the content of soluble sugar in kiwifruit and apple fruit (Miller et al., 1998; Wang et al., 2019). Drought treatment induces accumulation of flavonoids and anthocyanins in apple (Wang et al., 2020). Recently, transcriptome analysis of apricot fruit revealed that MYC and bHLH transcription factors may respond to stress and play a crucial role in flavor formation (Zhang et al., 2019). However, the regulatory mechanism of stress-mediated aroma accumulation remains unclear.

Apple (*Malus domestica*) is an economically important tree cultivated worldwide (Duan et al., 2017; Cornille et al., 2019). Ripening apple fruit produce approximately 350 volatile compounds, including aldehydes, alcohols, esters, ketones, and terpenes (Dimick and Hoskin, 1983; Song, 2007). Twenty types of volatile compounds are characteristic of the apple fruit aroma, which include *trans*-2-hexenal, hexanol, butyl acetate, hexyl acetate, and 2-methyl butyl acetate (Dixon and Hewett, 2000). With ripening of the fruit, the abundance of esters increases significantly (Rowan et al., 1999; Echeverrá et al., 2004). In ‘Golden Delicious’ apple, esters account for 80% of the total volatile aroma components (López et al., 2010). In ‘Golden Delicious’, 23 functional LOXs have been identified of which MdLOX1a and MdLOX5e might be involved in volatile component production (Vogt et al., 2013). *LOX* genes play a crucial role in the lipoxygenase pathway. However, little information is available on the regulation of LOXs in apple.

In this study, we selected a ripening-related gene, *MdLOX1a*, to investigate ester biosynthesis based on the results of a correlation analysis and overexpression of *MdLOX1a* in apple calli. A novel abiotic stress gene, *MdASG1*, was identified by yeast one-hybrid library screening. *MdASG1* responded to salt stress, directly bound to the promoter of *MdLOX1a* and activated its transcript, and, subsequently, enhanced the synthesis of aroma compounds. Overexpression of *MdASG1* in tomato fruit increased the production of volatile aroma compounds under salt stress. Overall, the findings provide new insights into the regulation of aroma compound production and a potential strategy to develop and utilize saline-alkali land to produce high-quality fruit, thereby reducing pressure on arable land and ensuring national food security.

## RESULTS

### *MdLOX1a* is involved in ester formation in apple and phylogenetic analysis of LOXs

We sampled apple fruit at four developmental and ripening stages (Fig. 1A) for gas chromatography–mass spectrometry (GC-MS) analysis. With progression of ripening, large amounts of esters were produced. In ripe fruit, the ester content attained about 14 μg·g^−1^ fresh weight, which was almost seven times that of immature fruit at 57 days after full bloom (DAFB) (Fig. 1B). The lipoxygenase pathway is one pathway for the synthesis of esters and the crucial participating enzyme is lipoxygenase. In apple, eight groups of LOXs are involved in the lipoxygenase pathway (Vogt et al., 2013). To clarify the key lipoxygenase genes in fruit ripening, we analyzed eight lipoxygenase genes from each group by quantitative real-time PCR analysis during fruit development and ripening (Fig. 1C, Supplemental Fig. S1). As the fruit matured, the transcript level of *MdLOX1a* increased significantly, consistent with the rate of ethylene release; in particular, *MdLOX1a* transcript abundance increased about 122-fold at the ripening stage compared with that of immature fruit (57 DAFB) (Fig. 1, C and D). A significant positive correlation was observed between the expression profile of *MdLOX1a* and ester content during apple fruit development and ripening (*r* = 0.989, *P* < 0.05) (Fig. 1E, Supplemental Fig. S2), which indicated that *MdLOX1a* may be a maturity-related gene. To further analyze the relationship between *MdLOX1a* and ester synthesis, we quantified *MdLOX1a* transcript levels (Fig. 1F), lipoxygenase activity (Fig. 1G), and ester content (Fig. 1H) in ripe fruit of eight popular apple cultivars (Supplemental Fig. S3). The transcript levels of *MdLOX1a* were positively correlated with lipoxygenase activity (*r* = 0.9464, *P* < 0.01; Fig. 1I). In addition, *MdLOX1a* transcript levels were positively correlated with ester content (*r* = 0.7408, *P* < 0.05) (Fig. 1J). These results indicated that *MdLOX1a* may be a critical gene involved in volatile ester biosynthesis.

**Figure 1.**
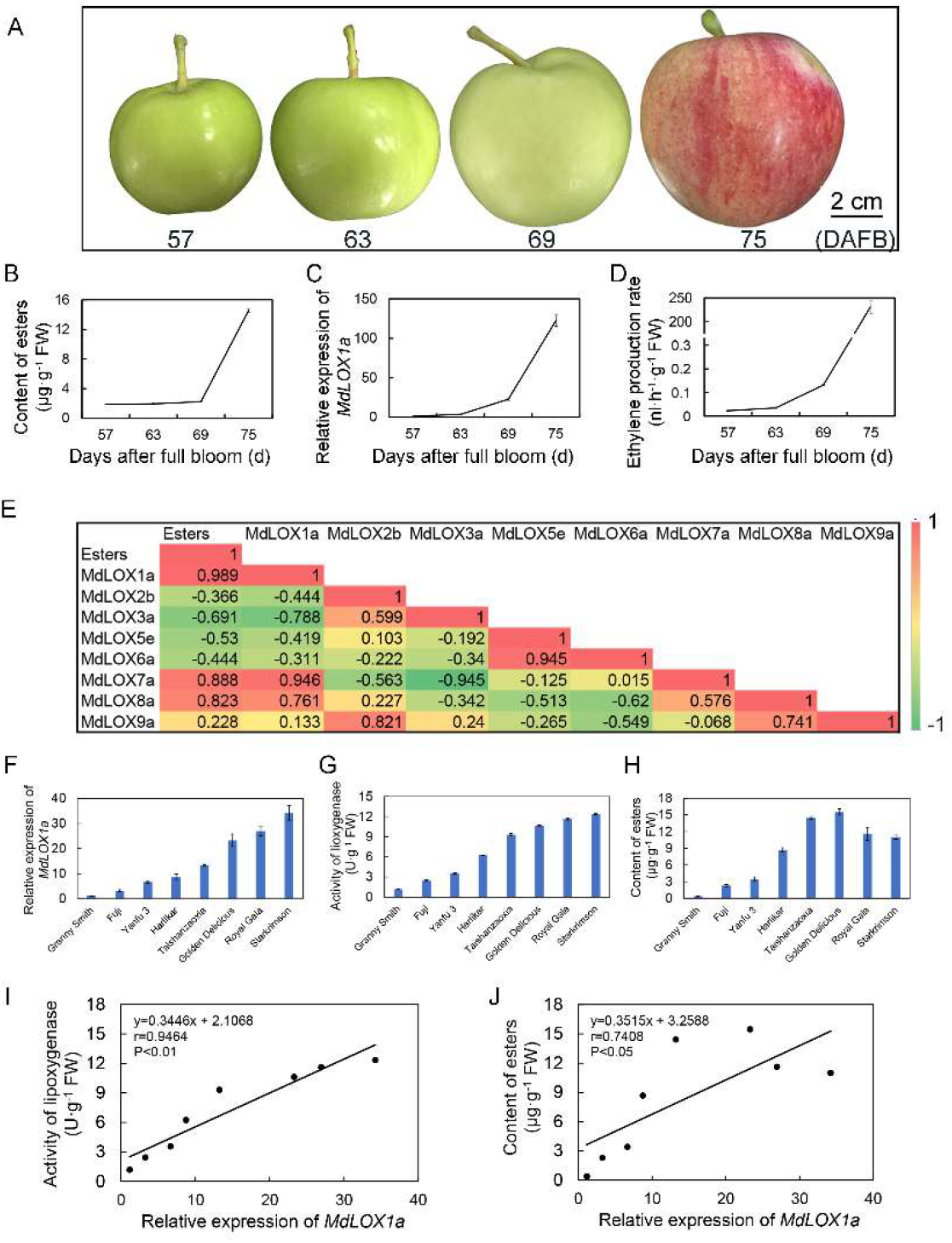
*MdLOX1a* is involved in ester formation in apple. A, Apple ‘Taishanzaoxia’ fruit were harvested at 57, 63, 69, and 75 days after full bloom (DAFB). Bar = 2 cm. B, Ester content during apple fruit development and ripening. C, Transcript level of *MdLOX1a* during apple fruit development and ripening quantified by qRT-PCR. *MdActin* was used as an internal control gene. D, Ethylene release rate during apple fruit development and ripening. E, Correlation analysis of *MdLOX* expression and ester content in apple fruit at the ripening stage. F, Relative expression of *MdLOX1a* in fruit of eight apple cultivars at the ripening stage. *MdActin* was used as an internal control gene. G, Lipoxygenase activity in fruit of eight apple cultivars at the ripening stage. H, Ester content in fruit of eight apple cultivars at the ripening stage. I, Linear regression analysis between *MdLOX1a* expression and lipoxygenase activity in fruit of eight apple cultivars. J, Linear regression analysis between *MdLOX1a* expression and ester content in fruit of eight apple cultivars. Error bars represent the standard deviation of three independent biological replicates. FW, Fresh weight. Significant differences were determined using Tukey one-way analysis of variance (ANOVA) with SPSS Statistics 22 (*P < 0.05 and **P < 0.01).

Functional LOXs have been identified in many plant species, including common bean (*Phaseolus vulgaris*), tomato, kiwifruit, Arabidopsis, grape, rice (*Oryza sativa*), persimmon (*Diospyros kaki*), and oriental melon (*Cucumis melo*) (Porta et al., 1999; Chen et al., 2004; Zhang et al., 2006; Bannenberg et al., 2009; Podolyan et al., 2010; Umate, 2011; Hou et al., 2015; Xing et al., 2020). Furthermore, 23 functional LOXs in the lipoxygenase pathway have been identified in the genome of ‘Golden Delicious’ apple (Vogt et al., 2013). In the present study, the amino acid sequence of 58 LOXs from 14 plant species was analyzed. The LOXs were mainly divided into two groups comprising 9-LOXs and 13-LOXs, respectively. *MdLOX1a* and *MdLXO7a*, which showed similar expression trends during apple fruit development and ripening, were grouped with 9-LOXs (Supplemental Fig. S4), which are classified as type 1 LOXs (Vogt et al., 2013).

### Overexpression of *MdLOX1a* increases fatty acid-derived volatile content in apple calli and subcellar localization of MdLOX1a

To analyze the function of *MdLOX1a* in volatile aroma biosynthesis, we generated *MdLOX1a*-overexpressing transgenic ‘Orin’ calli (Fig. 2, A and B). The content of fatty acid-derived volatiles, including esters, was significantly increased compared with that of the wild type (WT) (Fig. 2C). In addition, the corresponding synthetic genes in the lipoxygenase pathway were up-regulated (Fig. 2D). Taken together, these results indicated that *MdLOX1a* was associated with ester content. The construct 35S::MdLOX1a-GFP was generated to determine the subcellular localization of MdLOX1a. Strong green fluorescence signal was detected in the cytoplasm of tobacco (*Nicotiana benthamiana*) leaves (Fig. 2E), consistent with the subcellular localization predicted using Cell-PLoc 2.0.

**Figure 2.**
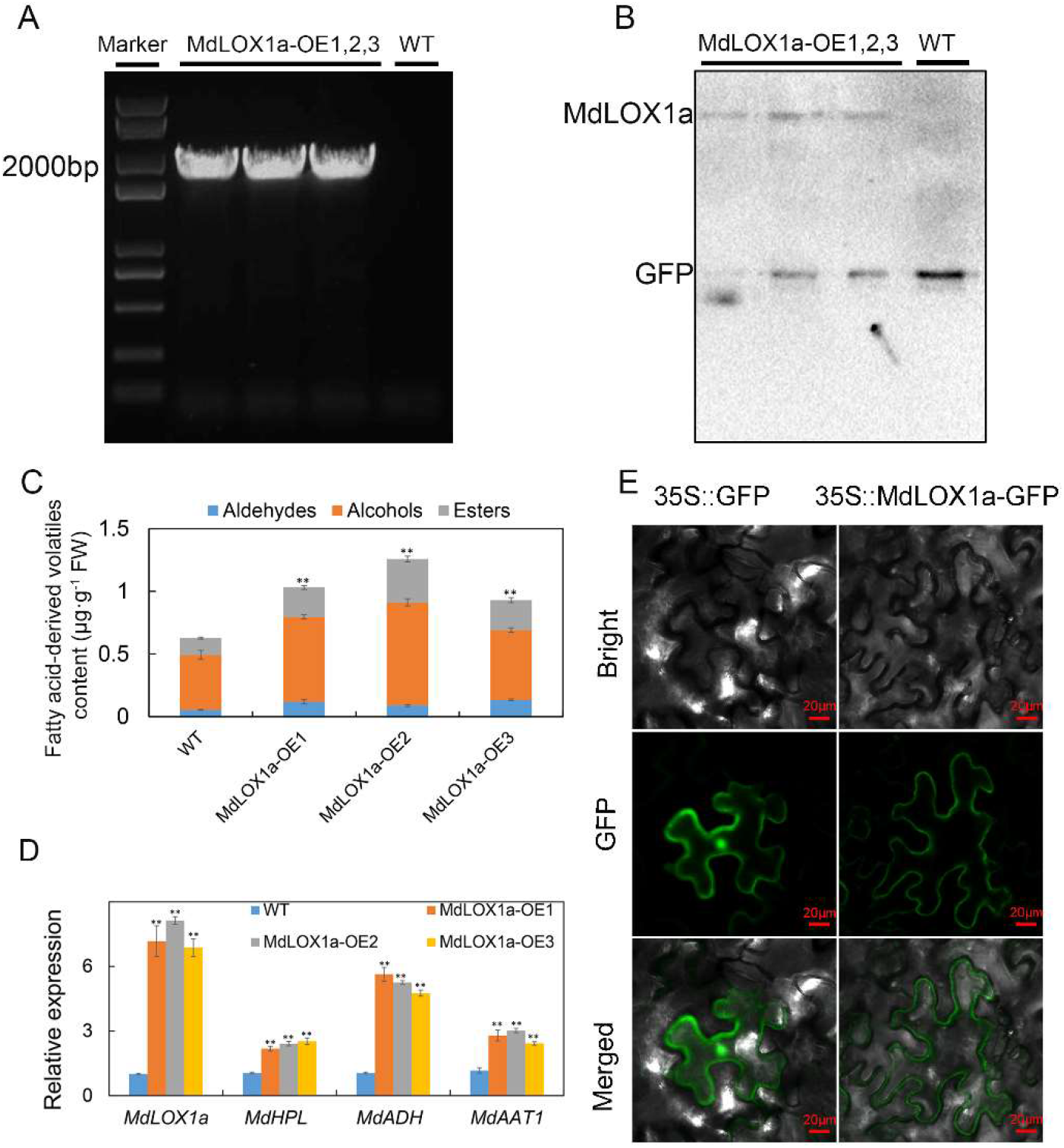
Overexpression of *MdLOX1a* increased fatty acid-derived volatile content in apple calli and subcellar localization of MdLOX1a. A and B, *MdLOX1a* overexpression in ‘Orin’ calli verified by PCR amplification (A) and western blotting (B). The 35S and MdLOX1a-PRI101-R primers were used for verification of transformants. C, Fatty acid-derived volatile contents in wild-type apple calli (WT) and *MdLOX1a*-overexpression apple calli (MdLOX1a-OE). D, Relative expression of fatty acid-derived volatile biosynthesis genes in WT and MdLOX1a-OE transgenic apple calli. *MdActin* was used as an internal control gene. E, Subcellular localization of MdLOX1a in tobacco leaves. MdLOX1a was mainly expressed in the cytoplasm of tobacco leaves. Bars = 20 μm. Error bars represent the standard deviation of three independent biological replicates. Asterisks indicate statistical significance (**P < 0.01 and *P < 0.05).

### A novel gene, *MdASG1*, is a direct regulator of the *MdLOX1a* promoter and activates its expression

Given that *MdLOX1a* is a crucial gene in volatile ester biosynthesis, we used the *MdLOX1a* promoter as bait to conduct yeast one-hybrid library screening. We identified a novel gene, designated *MdASG1* (accession number: XM_029093686), that was capable of binding to the promoter of *MdLOX1a* in the presence of 400 ng·mL^−1^ aureobasidin A (AbA) (Fig. 3A, Supplemental Fig. S5). The amino acid sequence of MdASG1 showed 70% and 73% similarity with Arabidopsis AtASG1 and potato (*Solanum commersonii*) ScASG1, respectively (Supplemental Fig. S6). The latter two genes both respond to stress treatment (Batelli et al., 2012). To determine the specific binding site of *MdASG1*, the promoter of *MdLOX1a* was divided into four fragments. Yeast one-hybrid assays showed that *MdASG1* bound to the *p4MdLOX1a* fragment (Fig. 3B).

**Figure 3.**
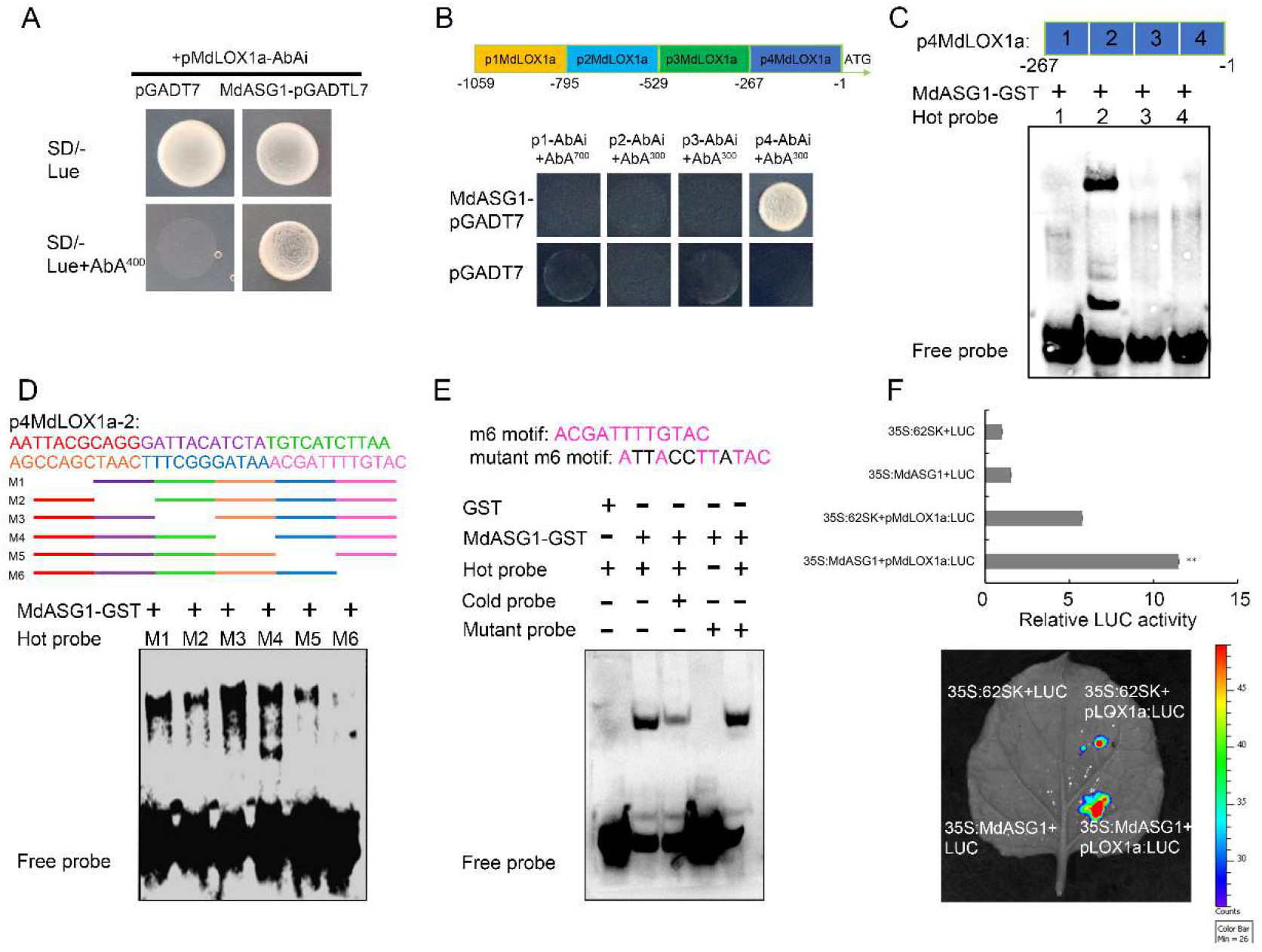
MdASG1 binds to the promoter of *MdLOX1a* and activates its expression. A, Yeast one-hybrid assays showing binding between MdASG1 and the promoter of *MdLOX1a*. B, Yeast one-hybrid assays showing binding between MdASG1 and the fourth segment of the promoter of *MdLOX1a* (*p4MdLOX1a*). C, Four segments of *p4MdLOX1a*. Electrophoretic mobility shift assay (EMSA) showing binding of MdASG1 to the −795~-529bp segment of *p4MdLOX1a* (*p4MdLOX1a-2*). D, Design of biotin-labeled probes (M1–M6) for partial deletion of the fragment *p4MdLOX1a-2*. The M6 fragment showed no binding with MdASG1. E, EMSA showing the binding of MdASG1 to the m6 motif in *MdLOX1a*. The hot probe was a biotin-labeled fragment. The cold probe was a nonlabeled fragment. The mutant probe contained five nucleotide mutations. The symbol + or – indicates the presence or absence of specific probes. F, Dual-luciferase assay verifying that MdASG1 transformation activated the *MdLOX1a* promoter. Error bars represent the standard deviation of three independent biological replicates. Asterisks indicate statistical significance (**P < 0.01 and *P < 0.05).

An electrophoretic mobility shift assay (EMSA) demonstrated that only −795~-529bp of *p4MdLOX1a* (*p4MdLOX1a-2*) contained a binding site (Fig. 3C). Partial deletion of the fragment *p4MdLOX1a-2* was performed to generate six individual fragments. Interestingly, binding was not observed in the absence of the m6 region (Fig. 3D). Therefore, we concluded that the specific binding motif of *MdASG1* was located in the m6 region. Addition of a cold probe weakened the binding. When the binding sites were mutated, binding was eliminated (Fig. 3E). A dual-luciferase reporter assay, performed to clarify the transcriptional regulation of *MdASG1* on *MdLOX1a*, indicated that MdASG1 targeted *MdLOX1a* as a transcriptional activator (Fig. 3F).

### Correlation of *MdASG1* expression with *MdLOX1a* transcript level and ester content

To further explore the relationship between *MdASG1* expression and aromatic compound synthesis, we analyzed the transcript level of *MdASG1* during apple fruit development and ripening (Fig. 4A), which was consistent with the changes in ester content. Subsequently, the expression profile of *MdASG1* among eight apple cultivars was examined (Fig. 4B). Correlation analysis among the cultivars revealed that *MdASG1* expression was positively correlated with *MdLOX1a* expression (*r* = 0.7690, *P* < 0.05) (Fig. 4C). Furthermore, the expression profile of *MdASG1* was correlated with ester content among the cultivars (*r* = 0.8207, *P* < 0.05) (Fig. 4D). Taken together, these results suggested that *MdASG1* was a candidate gene involved in the lipoxygenase biosynthesis pathway. Subcellular localization showed that *MdASG1* was uniformly distributed in all subcellular compartments (Fig. 4E).

**Figure 4.**
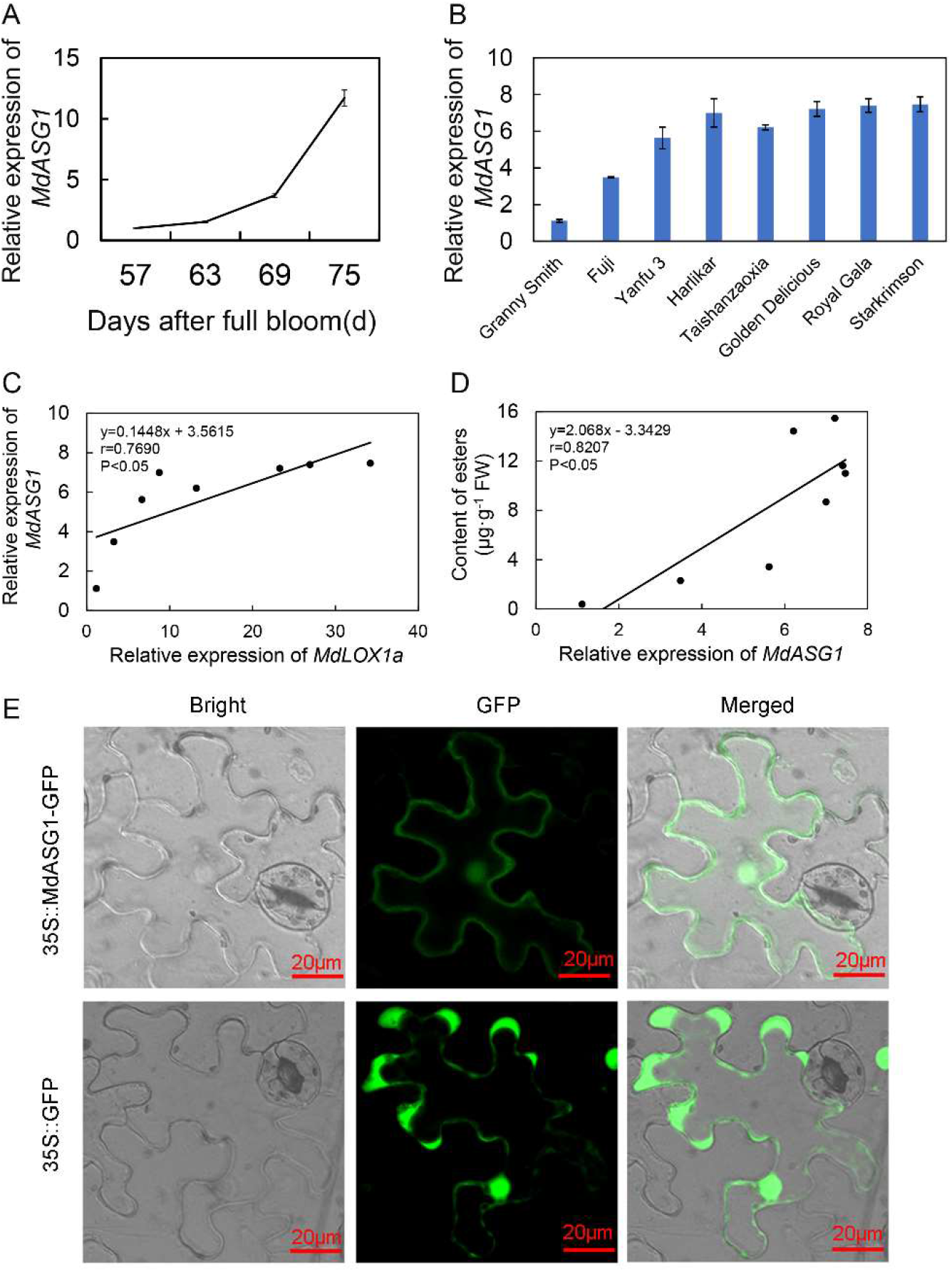
*MdASG1* is involved in ester biosynthesis in apple. A and B, Transcript levels of *MdASG1* during apple fruit development and ripening (A), and in fruit of eight popular apple cultivars at ripening (B). *MdActin* was used as an internal control gene. C, Linear regression analysis between *MdLOX1a* expression and *MdASG1* expression in fruit of eight apple cultivars. D, Linear regression analysis between *MdASG1* expression and ester content in fruit of eight apple cultivars. E, Subcellular localization of MdASG1 in tobacco leaves. Bars = 20 μm. Error bars represent the standard deviation of three independent biological replicates. Significant differences were determined using Tukey one-way analysis of variance (ANOVA) with SPSS Statistics 22 (*P < 0.05 and **P < 0.01).

### Changes in fatty acid-derived volatile content caused by transient overexpression of *MdASG1* or silencing of *MdASG1* in apple

Given the positive correlation between the expression of *MdASG1* and *MdLOX1a*, as well as the ester content (Fig. 4), we hypothesized that *MdASG1* plays a role in regulating aroma compound biosynthesis. To test this hypothesis, we transiently overexpressed *MdASG1* in ‘Yinv’ apple by injecting *Agrobacterium tumefaciens* infiltration buffer containing the target gene or the empty vector (Fig. 5A). An approximately 2-fold increase in *MdASG1* transcript levels and then a about 6-fold increase in *MdLOX1a* transcript levels were observed (Fig. 5B). These changes were accompanied by higher contents of fatty acid-derived volatiles, including 1-hexanol, hexyl acetate, and 2-hexen-1-ol, acetate, (Z), compared with transient expression of the empty vector 35S::GFP (Fig. 5, C–E). In addition, we transiently silenced *MdASG1* (Fig. 5F). The opposite results were observed in *MdASG1*-silenced fruits and fatty acid-derived volatile contents were significantly inhibited at the TRV-MdASG1 injection sites (Fig. 5G).

**Figure 5.**
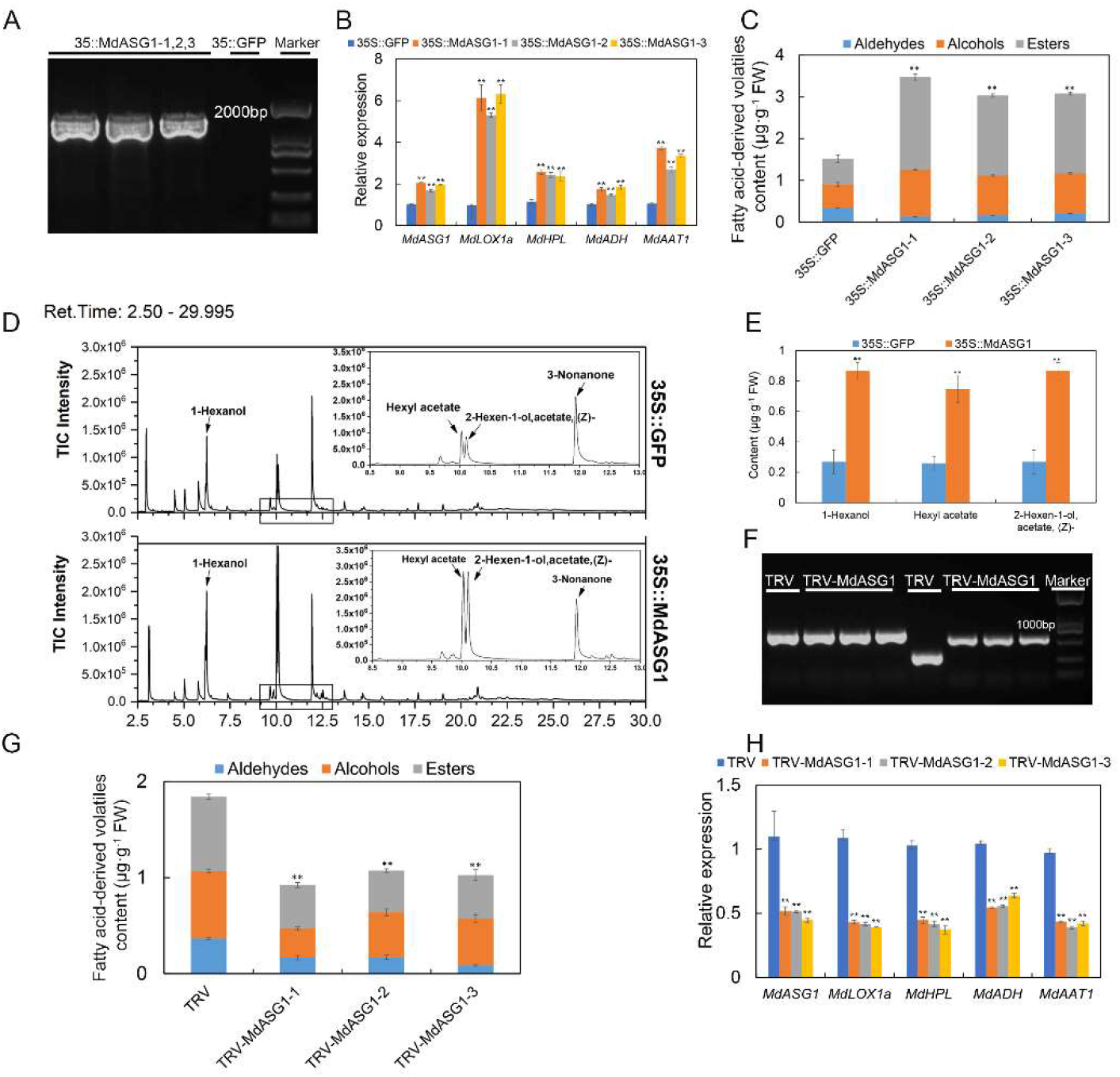
Transient overexpression or silencing of *MdASG1* in apple fruit. A, Transient overexpression of *MdASG1* was confirmed by PCR amplification. The GFP-F and MdASG-PHB-R primers were used for verification of transformants. B, Relative expression of *MdASG1* and fatty acid-derived volatile biosynthesis genes in apple fruit with transient overexpression of *MdASG1* (35::MdASG1) and the empty vector (35S::GFP). *MdActin* was used as an internal control gene. C, Fatty acid-derived volatile content in 35S::GFP and 35::MdASG1 transgenic apple fruit. D, Mass spectra of 35S::GFP and 35::MdASG1 transgenic apple fruit. E, Contents of 1-hexanol, hexyl acetate, and 2-hexen-1-ol, acetate, (Z) in 35S::GFP and 35::MdASG1 transgenic apple fruit. F, Transient silencing of *MdASG1* was confirmed by PCR amplification. The TRV1-F and TRV1-R primers were used in lanes 1–4 from the left, and the TRV2-F and TRV2-R primers were used in lanes 5–8 from the left. G, Fatty acid-derived volatiles content in apple fruit with transient silencing of *MdASG1* (TRV-MdASG1) and the empty vector (TRV). H, Relative expression of *MdASG1* and fatty acid-derived volatile biosynthesis genes in TRV and TRV-MdASG1 transgenic apple fruit. *MdActin* was used as an internal control gene. Error bars represent the standard deviation of three independent biological replicates. Asterisks indicate statistical significance (**P < 0.01 and *P < 0.05).

Silencing of *MdASG1* led to a corresponding decrease in the transcript level of genes associated with the lipoxygenase pathway (Fig. 5H).

### Changes in fatty acid-derived volatile content caused by stable overexpression of *MdASG1*

To provide further evidence of *MdASG1*-mediated fatty acid-derived volatile production, we generated *MdASG1*-overexpression ‘Orin’ calli (Fig. 6, A and B).

**Figure 6.**
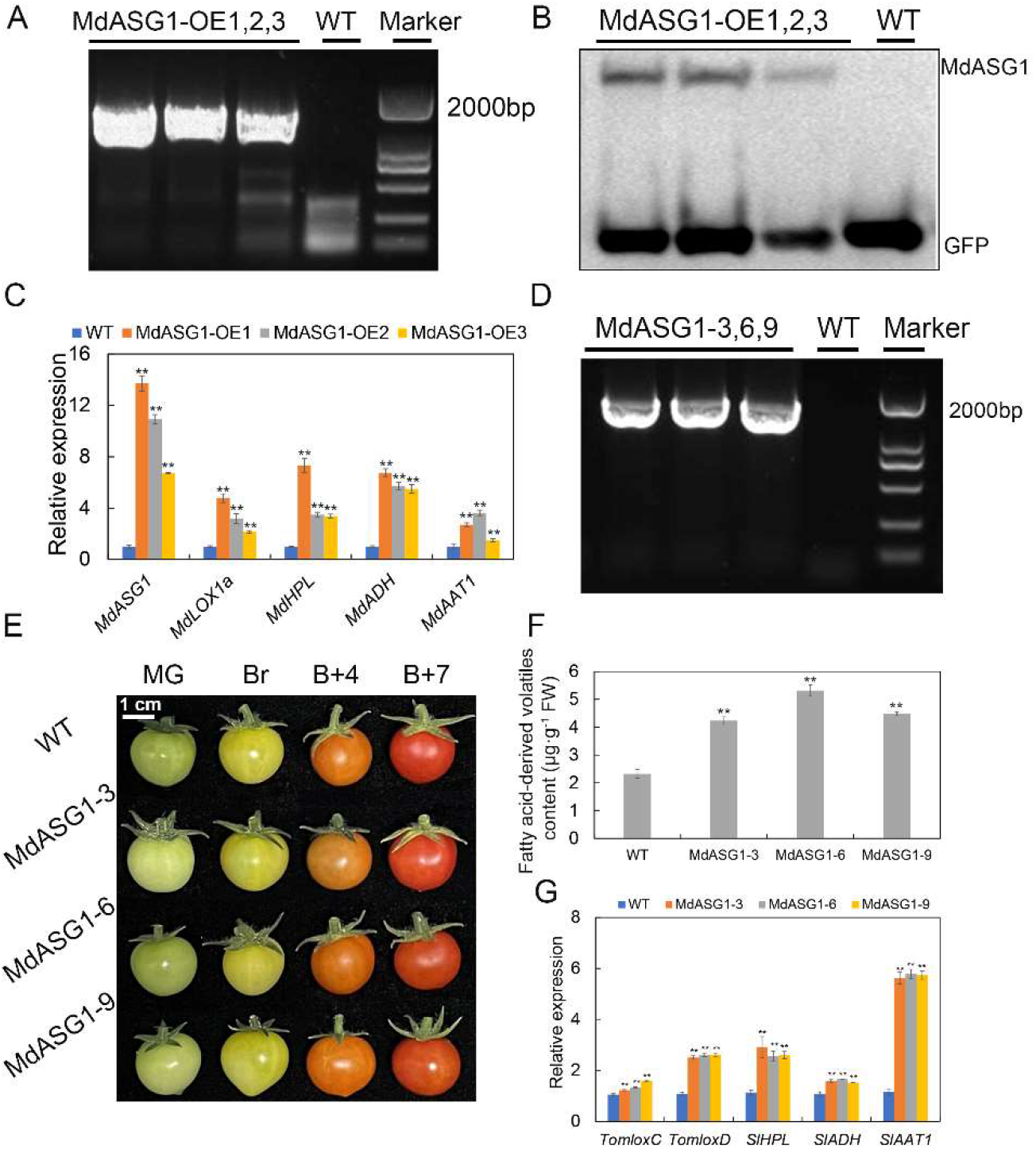
Stable overexpression of *MdASG1* in apple and tomato fruit. A and B, *MdASG1* overexpression in apple ‘Orin’ calli verified by PCR amplification (A) and western blotting (B). The 35S and MdASG1-PRI101-R primers were used for verification of transformants. C, Relative expression of *MdASG1* and fatty acid-derived volatile biosynthesis genes in *MdASG1*-overexpressing ‘Orin’ (MdASG1-OE) and wild-type (WT) calli. *MdActin* was used as an internal control gene. D, *MdASG1* overexpression in tomato verified by PCR amplification. The 188F and MdASG1-PCB302-R primers were used for verification of transformants. E, Fruit of tomato ‘Micro-Tom’ overexpressing *MdASG1*. F, Fatty acid-derived volatile content in fruit of wild-type Micro-Tom (WT) and *MdASG1*-overexpression tomato (MdASG1-3,6,9). G, Relative expression of fatty acid-derived volatile biosynthesis genes in fruit of WT and MdASG1-3,6,9 transgenic tomato. *SlActin* was used as an internal control gene. Error bars represent the standard deviation of three independent biological replicates. Asterisks indicate statistical significance (\*\**P* < 0.01 and \**P* < 0.05).

Overexpression of *MdASG1* caused upregulation in *MdLOX1a* expression and that of other genes in the lipoxygenase pathway compared with the control calli (WT) (Fig. 6C). To rapidly generate transgenic fruit, we overexpressed *MdASG1* in tomato ‘Micro-Tom’ and obtained the lines MdASG1-3, MdASG1-6, and MdASG1-9 (Fig. 6, D and E). Ripening fruit of these overexpression lines accumulated higher contents of volatiles than the WT (Fig. 6F). The transcript levels of the corresponding synthase genes involved in the lipoxygenase pathway in transgenic tomato fruit were significantly higher than those of WT tomato (Fig. 6G). To summarize, these results suggested that *MdASG1* promotes fatty acid-derived volatile biosynthesis by activating the transcript of *MdLOX1a* in the lipoxygenase pathway.

### Overexpression of *MdASG1* confers enhanced salt tolerance and accumulation of higher contents of fatty acid-derived volatiles under salt treatment

*MdASG1* showed high homology with *AtASG1*. Therefore, we speculated that *MdASG1* may respond to abiotic stress similar to *AtASG1*. As expected, *MdASG1* transcript levels were higher in response to NaCl treatment in tissue-cultured plantlets of ‘Royal Gala’ (Fig. 7A) and ‘Orin’ calli (Fig. 7B), and especially in *MdASG1*-overexpression ‘Orin’ calli (Fig. 7B). Similarly, *MdASG1*-overexpression ‘Orin’ calli were more tolerant to salt stress than the control (Fig. 7C) and the transcription of stress-related genes was up-regulated (Supplemental Fig. S7). Interestingly, the transcript levels of genes in the lipoxygenase pathway were up-regulated in response to 50 mM NaCl treatment for 20 d, especially in calli overexpressing *MdASG1* (Fig. 7D). Similar results were observed in tomato; the transcript levels of the tomato homolog *SlASG1* (Fig. 7E) and genes in the lipoxygenase pathway were up-regulated with 200 mM NaCl treatment in WT and transgenic tomato fruit (Supplemental Fig. S8). The contents of fatty acid-derived volatiles increased accordingly under the salt treatment in WT and transgenic tomato fruit (Fig. 7F). The transgenic tomato plants exhibited a significant increase in tolerance to salt stress (Fig. 7G), higher photosynthesis capacity (Fig. 7H), and reduced oxidative stress (Fig. 7I) compared with the WT.

**Figure 7.**
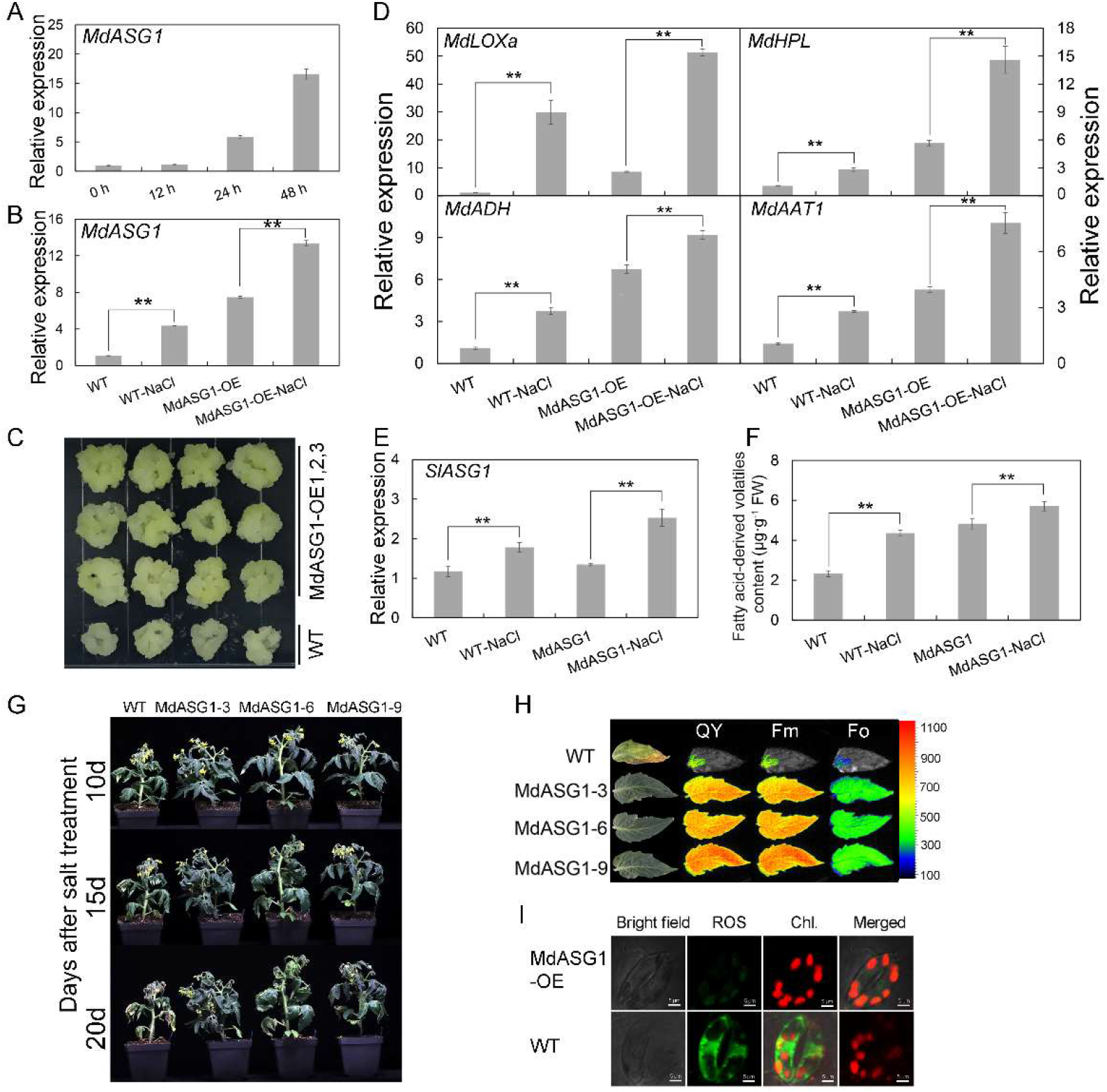
*MdASG1* enhances plant salt tolerance and mediates enhanced accumulation of fatty acid-derived volatiles under salt stress. A, Relative expression of *MdASG1* in wild-type tissue-cultured apple plantlets under 200 mM NaCl treatment. *MdActin* was used as an internal control gene. B, Transcriptional changes in *MdASG1* in response to 50 mM NaCl treatment in wild-type ‘Orin’ calli (WT) and *MdASG1*-overexpressing transgenic lines (MdASG1-OE). *MdActin* was used as an internal control gene. C, WT and MdASG1-OE transgenic ‘Orin’ calli treated with 50 mM NaCl. D, Transcriptional changes in fatty acid-derived volatile biosynthesis genes under 50 mM NaCl treatment in WT and MdASG1-OE transgenic ‘Orin’ calli. *MdActin* was used as an internal control gene. E, Transcriptional changes in *SlASG1* under 200 mM NaCl treatment in ripening fruit of WT and *MdASG1*-overexpression (MdASG1) tomato. *SlActin* was used as an internal control gene. F, Changes in fatty acid-derived volatile content under 200 mM NaCl treatment in ripening fruit of WT and MdASG1 transgenic tomato. G, Phenotype of WT and MdASG1-3,6,9 transgenic tomato plants subjected to 200 mM NaCl treatment for 10, 15, and 20 d. H, Chlorophyll fluorescence in tomato leaves after NaCl treatment for 20 d. I, Fluorescence of reactive oxygen species in tomato leaf cells after NaCl treatment for 20 d. Bars = 5 μm. Error bars represent the standard deviation of three independent biological replicates. Asterisks indicate statistical significance (**P < 0.01 and *P < 0.05).

**Figure 8.**
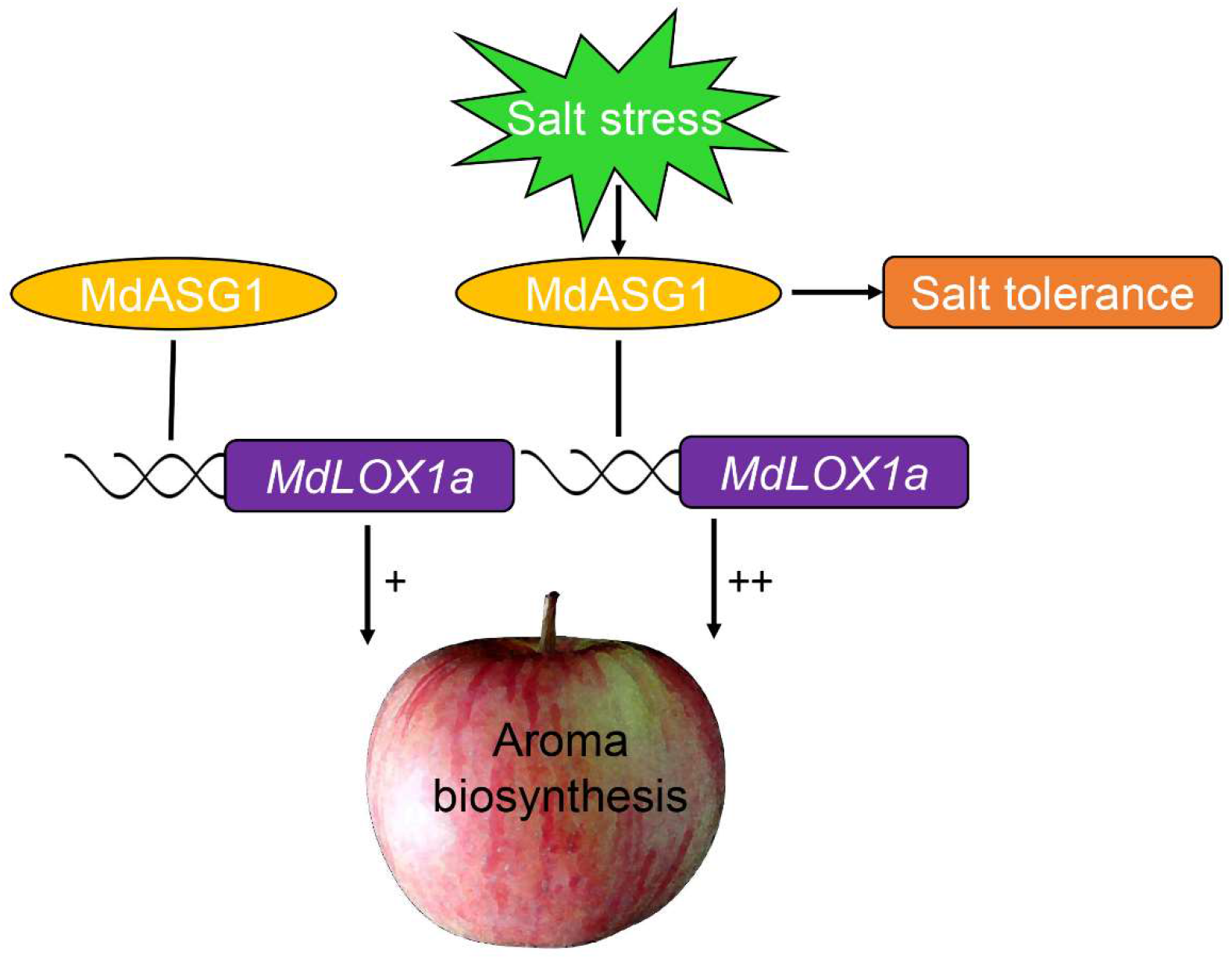
Proposed model for MdASG1 modulation of aroma compound biosynthesis in apple. MdASG1 can increase aroma compound biosynthesis by binding to the promoter of *MdLOX1a*. Under moderate salt stress, MdASG1 enhances tolerance to salt stress and promotes accumulation of aroma compounds in fruit.

The expression of stress-related genes was up-regulated in transgenic tomato plants (Supplemental Fig. S9). In summary, these results suggest that *ASG1* is involved in the synthesis of volatile aroma compounds in apple and tomato, and higher contents of aroma compounds accumulate under salt stress.

## DISCUSSION

Fruit flavor comprises a complex set of interactions between taste and aroma (Brückner and Wyllie, 2008). Aroma is a mixture of various volatile compounds and is an important quality trait that influences consumer acceptance. The synthesis and accumulation of aroma compounds are increased in ripening apple fruit of which esters account for 80% of the volatiles (López et al., 1998; Lavilla et al., 1999; López et al., 2010). Lipoxygenase is an important contributor to fruit ester production. In pepino fruit during ripening, three LOX genes responsible for aroma compound biosynthesis, namely *SmLOXD*, *SmLOXB*, and *SmLOX5-like2*, are up-regulated (Contreras et al., 2017). In kiwifruit, *AdLox1* and *AdLox5* are up-regulated during ripening and are involved in fruity aroma ester synthesis (Zhang et al., 2009). *MdLOX1a* is associated with a quantitative trait locus for volatile esters in apple (Schiller et al., 2015). However, the specific function of *MdLOX1a* in ester synthesis requires further study. In the present study, *MdLOX1a* and *MdLOX7a* were up-regulated during fruit ripening, consistent with results reported by Schiller et al. (2015). We determined that *MdLOX1a* is involved in ester biosynthesis based on the significant positive correlations between *MdLOX1a* expression and ester content. In addition, *MdLOX1a* overexpression in apple ‘Orin’ calli increased the ester content. Therefore, we speculated that *MdLOX1a* is a crucial gene in the lipoxygenase pathway. Plant LOXs are localized in the cytoplasm or chloroplasts. Tomato TomloxC is involved in the synthesis of C6 flavor compounds, which are localised in the chloroplasts (Chen et al., 2004). MdLOX1a is localized in the cytoplasm to participate in ester synthesis. Similarly, in kiwifruit, AdLox5 participates in fruity aroma ester synthesis in the cytoplasm (Zhang et al., 2006; Zhang et al., 2009). Phylogenetic analysis revealed that MdLOX1a can be classified as a 9-LOX. However, MdLOX1 is reported to have a dual positional specific function generating 9- and 13-hydroperoxides (Schiller et al., 2015).

Transcriptional regulation of fruit aroma components has been widely reported in plants. However, previous research has mainly focused on terpene biosynthesis. For instance, multiple transcription factors of the MYC2, NAC, EIL, AP2/ERF, and MYB families (Hong et al., 2012; Li et al., 2015; Nieuwenhuizen et al., 2015; Shen et al., 2016; Li et al., 2017; Jian et al., 2019) are involved in terpene synthesis by directly activating the terpene synthase *TPS*. Recently, transcription factors of the bZIP, NAC, and Dof families have been reported to play important roles in ester biosynthesis by regulating expression of the structural gene *AAT* in the lipoxygenase pathway (Guo et al., 2018; Zhang et al., 2020; Cao et al., 2021; Wang et al., 2022). *LOX* is a crucial structural gene in the lipoxygenase pathway, but the regulation of *LOX* is rarely reported. Given the observation that *MdLOX1a* mediates fruit ester biosynthesis, we used *MdLOX1a* as a candidate gene and identified a novel abiotic stress gene, *MdASG1*, which activated *MdLOX1a* expression by directly binding to its promoter. Furthermore, overexpression of *MdASG1* in apple fruit increased aroma compound production, whereas synthesis of these compounds was decreased by *MdASG1* silencing. In *Saccharomyces cerevisiae*, the zinc cluster transcriptional regulator Asg1 is an activator of stress-responsive genes, which involves fatty acid utilization (Jansuriyakul et al., 2016). However, MdASG1 and Asg1 of *S. cerevisiae* are entirely unrelated proteins.

The function of ASG (ScASG1 and AtASG1) was first identified in *Solanum tuberosum* and *Arabidopsis thaliana*, and is a positive regulator of stress responses via an ABA-dependent pathway (Batelli et al., 2012). Amino acid sequence analysis revealed that MdASG1 showed high homology with Arabidopsis AtASG1 and potato ScASG1. In the current study, we observed a novel function for *ASG* in apple in mediating aroma compound biosynthesis. In addition, we observed that *MdASG1* performed similar functions to those of *ScASG1* and *AtASG1* in response to NaCl treatment (Batelli et al., 2012). Transgenic apple calli and tomato plants (MdASG1-3,6,9) exhibited significantly enhanced tolerance to salt stress, and higher photosynthesis capacity and lower oxidative stress in transgenic tomato plants compared with the WT. We cloned *MdASG1* into the PHB vector and observed that MdASG1 was uniformly distributed in all subcellular compartments. In contrast, potato ScASG1 is localized to the plasma membrane (Batelli et al., 2012). Overexpression of *DkLOX3* and *CaLOX1* in Arabidopsis plant confer increased tolerance to high salinity and drought stress by modulating stress-related genes and reactive oxygen species production (Hou et al., 2015; Lim et al., 2015). In oriental melon, *CmLOX10* positively regulates drought tolerance through JA-mediated stomatal closure (Xing et al., 2020). In tomato, overexpression of ω-3 fatty acid desaturases (FAD), which catalyze the conversion of linoleic acid (18:2) to linolenic acid (18:3) in the lipoxygenase pathway, enhance tolerance to cold stress (Dominguez et al., 2010). Therefore, we speculated that *MdASG1* might function by mediating the lipoxygenase pathway in the response to abiotic stress.

Abiotic stress strongly affects plant growth. However, the observation that moderate stress may improve fruit quality is usually overlooked. Some previous studies have examined stress-mediated fruit quality but were mainly focused on sweetness and anthocyanin production, and less frequently on fruit aroma. For example, mild salt stress improves strawberry fruit quality by increased accumulation of sucrose and the antioxidant compounds anthocyanins and catechins (Casierra-Posada and Riaño, 2006; Keutgen and Pawelzik, 2007; Galli et al., 2016). Similarly, in tomato, NaCl treatment increases the concentration of soluble solids not only as a result of reduction in water transport (Sato et al., 2006; Saito et al., 2008; Johkan et al., 2014). In grape, moderate salinity increases anthocyanin and soluble solid contents, but decreases aroma quality (Li et al., 2013). Conversely, in the present study, moderate salt stress increased the expression of lipoxygenase pathway-related genes in apple calli and tomato fruit, accompanied by increased accumulation of aroma compounds. Especially in *MdASG1*-overexpressing apple calli and tomato, *MdASG1* further improved fatty acid-derived volatile content under moderate salt stress. At the same time, tomato *SlASG1*, which is a homolog of apple *MdASG1*, was significantly up-regulated under moderate salt stress, accompanied by the increase in aroma compound synthesis. These results collectively indicate that *ASG1* is involved in salt-induced aroma biosynthesis via enhanced expression of genes in the lipoxygenase pathway.

The present results provide a theoretical foundation for exploitation of moderate salt stress to improve fruit quality, and may enable the prudent development and utilization of saline-alkali land to produce high-quality fruit. Rice and wheat (*Triticum aestivum* L.) are major crops grown worldwide, but their growth and yield are frequently restrained by salinity stress (Castro-Llanos et al., 2019; Yan et al., 2020). It is estimated that at least 20% of all irrigated lands are impacted by salinity stress (Pitman and Läuchli, 2002). Given that irrigated lands may be adversely affected by salinity, our findings may contribute to improved utilization of saline-alkali land for crop production. Ultimately, this would relieve pressure on arable land, thereby ensuring global food security.

In summary, in this study we observed that the lipoxygenase gene *MdLOX1a* is a crucial gene involved in ester biosynthesis. We identified an abiotic stress gene, *MdASG1*, that directly bound to the promoter of *MdLOX1a* and activated its transcription, and thus participated in the biosynthesis of fatty acid-derived volatiles in apple fruit. In addition, *MdASG1* expression was enhanced by NaCl stress. Transgenic apple calli and tomato plants (MdASG1-3,6,9) were more tolerant of salt stress than the WT, and transcript levels of genes in the lipoxygenase pathway were higher under salt stress compared with the non-stress condition, which may explain how moderate stress improves fruit quality. The present results provide insight into the regulatory mechanism by which *MdASG1* directly activates *MdLOX1a* expression to promote accumulation of aroma volatiles especially under moderate salt stress. Our findings establish a theoretical strategy for production of improved-quality apple fruit on moderately saline soil to meet the needs of consumers.

## MATERIALS AND METHODS

### Plant material and culture conditions

Apple ‘Taishanzaoxia’ fruit were harvested at 57, 63, 69, and 75 DAFB from Liaocheng, Shandong province, China. Fruit of eight apple cultivars were sampled at the ripening stage from Liaocheng, Shandong province, China. The ‘Orin’ apple calli used for genetic transformation were cultured on Murashige and Skoog (MS) medium supplemented with 0.8 mg·L^−1^ 6-benzylaminopurine (6-BA) and 1.5 mg·L^−1^ 2,4-dichlorophenoxyacetic acid (2,4-D) in the dark at 24℃. Tissue-cultured plantlets of ‘Royal Gala’ were subcultured on MS medium supplemented with 0.2 mg·L^−1^ indoleacetic acid and 0.5 mg·L^−1^ 6-BA under a 16 h/8 h (light/dark) photoperiod at 24℃. Tomato ‘Micro-Tom’ plants were grown in a greenhouse at 24℃ under a 16 h/8 h (light/dark) photoperiod. Fruit of ‘Yinv’ apple used in the transient transformation assays were harvested before coloring from trees in the germplasm nursery of the Shandong Institute of Pomology. Tobacco (*Nicotiana benthamiana*) plants used for subcellular localization and dual-luciferase assays were grown in a plant growth chamber at 24℃ under a 16 h/8 h (light/dark) photoperiod.

### Stress treatment

The shoot tip of 25-day-old ‘Royal Gala’ tissue-cultured plantlets was excised and transferred to liquid MS medium supplemented with 200 mM NaCl. After 12, 24, and 48 h treatment, sampled shoots were immediately frozen in liquid nitrogen and stored at −80℃ until use. Transgenic (MdASG1-OE) and control (WT) calli of uniform growth status were cultured on MS medium supplemented with 50 μM NaCl for salt stress treatment for 20 d. Tomato plants grown in a square plastic pot (10 cm diameter at the top, 7.5 cm diameter at the bottom, and 8.5 cm in height) were well watered before salt treatment. One-month-old tomato plants (WT and T_3_ transformants) of uniform growth were watered with 200 mM NaCl solution at 4-day intervals until the fruit were ripe. After 20 d salt treatment, the leaves were sampled for observation of chlorophyll fluorescence, ROS, and RNA extraction. The ripe tomato fruit were collected for GC-MS analysis. All of the above-mentioned treatments were applied with three biological replicates.

### Volatile collection and GC-MS analysis

Headspace solid-phase microextraction was used to collect fruit volatile compounds following the method of Lu et al. (2021). Fresh fruit tissue (5 g) cut into pieces or 5 g apple calli was transferred to a 50 mL conical flask, then 10 μL 3-nonanone (0.4 mg·mL^−1^) was added as an internal standard, sealed, and extracted at 45℃ for 40 min. Tomato fruit required an additional 10 mL saturated NaCl to extract volatile compounds. The SPME fiber coated with a 50/30 μm layer of divinylbenzene/carboxen/polydimethylsiloxane (DVB/CAR/PDMS) (Supelco, Bellefonte, PA, USA) was used for volatile collection. The gas chromatograph–mass spectrometer (GCMS-QP2010, Shimadzu, Kyoto, Japan) was equipped with a Rtx-5MS capillary column (30 m × 0.25 mm i.d. × 0.25 μm film thickness) (Restek Co., Bellefonte, PA, USA). High-purity helium was employed as the carrier gas with a constant flow rate of 2 mL·min^−1^. The GC started at 35℃ for 2 min, increased to 120℃ at 6℃·min^−1^, then increased to 180℃ at 10℃·min^-1^, and finally increased to 250℃ at 20℃·min^−1^ for 5 min. The transfer, MS source, and interface temperature were 250℃, 200℃, and 230℃, respectively. The mass spectra were acquired with 70 eV electron ionization energy. The volatile compounds were identified by matching with the NIST 2017 mass spectral library and compared with the linear retention index (LRI) values. The relative content of a volatile compound was determined using the peak area of the internal standard as a reference according to the total ion chromatogram (TIC). For each sample three biological replicates were analyzed.

### RNA extraction and quantitative real-time PCR

Total RNA was extracted from plant tissue using the FastPure^®^ Plant Total RNA Isolation Kit (Vazyme, Nanjing, China). The synthesis of cDNA was performed using HiScript^®^ II Reverse Transcriptase (Vazyme). The qRT-PCR analysis was performed with the ChamQ SYBR qPCR Master Mix (Vazyme) using a CFX Connect instrument (Bio-Rad, Hercules, CA, USA). The *MdActin* gene was used as an internal control for apple and the *SlActin* gene was used as an internal control for tomato. Gene relative expression analysis used the 2^−∆∆*C*t^ method (Livak and Schmittgen, 2001). The primers used for qRT-PCR in this study are listed in Supplemental Table S1.

### Determination of lipoxygenase activity

Lipoxygenase activity was determined using a lipoxygenase assay kit (mlbio, Shanghai, China). Briefly, fruit tissue was ground into powder in liquid nitrogen. The powder (0.1 g) was shaken and resuspended in 1 mL extract buffer, and then centrifuged for 20 min at 16000 ×*g* at 4℃. The supernatant was the enzyme extract. The determination was performed by adding 20 μL enzyme, 160 μL buffer solution reagent, and 20 μL substrate solution. Lipoxygenase activity was measured as the increase in absorbance at 234 nm over 1 minute. One unit of enzyme activity was defined as the change in absorbance of 0.01 at 25℃ per minute per gram of tissue. Each sample was analyzed with three biological replicates.

### Phylogenetic analysis

A phylogenetic tree was constructed from a multiple alignment of 58 LOX amino acid sequences from 14 plant species. The sequences were downloaded from the Genome Database for Rosaceae (https://www.rosaceae.org/) or the National Center for Biotechnology Information database (http://www.ncbi.nlm.nih.gov/). The phylogenetic tree was constructed using the neighbor-joining method with MEGA X. A bootstrap analysis with 1000 replicates was performed to evaluate the reliability of the tree topology. The accession numbers of the LOX sequences are listed in Supplemental Table S4.

### Subcellular localization of MdLOX1a and MdASG1

The full-length coding sequence (CDS) of *MdLOX1a* or *MdASG1* was inserted into the pHB vector carrying the 35S promoter and then introduced into *A. tumefaciens* strain GV3101 using the freeze–thaw method. *Agrobacterium* containing the target gene was resuspended in infiltration buffer (10 mM MES, 10 mM MgCl_2_, and 150 μM acetosyringone, pH 5.5–5.7) and injected into 1-month-old tobacco leaves. The fluorescence signal was imaged after infiltration for 2 d using a confocal laser microscope (LSM880, Carl Zeiss, Oberkochen, Germany). The primers used for vector construction are listed in Supplemental Table S2.

### Yeast one-hybrid assay

To screen for proteins that potentially bind to the promoter of *MdLOX1a*, we used the Matchmaker^®^ Gold Yeast One-Hybrid Library Screening System (Clontech, Mountain View, CA, USA) following the manufacturer’s instructions. The *MdLOX1a* promoter (fragment length 1059 bp) was cloned into the pAbAi vector and the linearized plasmid was transformed into the yeast strain Y1H Gold. The optimal AbA screening concentration was determined in accordance with the manufacturer’s instructions. Total RNA extracted from apple ‘Taishanzaoxia’ fruit at different developmental stages was used to construct the prey cDNA library. The library plasmid (10 μL) was transformed into the MdLOX1a-pAbAi Y1H Gold strain to screen the novel protein. In addition, the identified protein MdASG1 was cloned into the pGADT7 vector to confirm the result. The promoter of *MdLOX1a* was divided into four fragments (*p1MdLOX1a* to *p4MdLOX1a*) to identify the binding site. The corresponding primers used for vector construction are listed in Supplemental Table S2.

### Dual-luciferase reporter assay

The full-length CDS of *MdASG1* was inserted into the pGreenⅡ62-SK vector. The *MdLOX1a* promoter was inserted into the pGreenⅡ0800-Luc vector. The recombinant plasmids were expressed transiently in tobacco leaves by *A. tumefaciens*-mediated genetic transformation using the same method as described for the subcellar localization assay. The In Vivo Imaging System (Xenogen, Alameda, CA, US) was used to detect luminescence. The luciferase activities were measured using the Dual-Luciferase^®^ Reporter Assay System (Promega, Madison, WI, US).

### EMSA

The EMSA was conducted using the Lightshift Chemiluminescent EMSA kit (Thermo, New York, NY, USA) in accordance with the manufacturer’s instructions as described previously by Zhang et al. (2018). The full length CDS of *MdASG1* was inserted into the pGEX-4T vector. The constructed vector was introduced into *Escherichia coli* strain BL21 to induce protein production, and then was purified using the GST-tag Protein Purification Kit (Beyotime, Shanghai, China) following the manufacturer’s instructions. The biotin-labeled probe and MdASG1-GST protein were mixed in the binding buffer and incubated at 24℃ for 15 min. The GST protein was used as the control. The unlabeled probes were used as competitors. The probes used in the EMSA assay are listed in Supplemental Table S3.

### Fluorescence detection of reactive oxygen species

Reactive oxygen species were detected with fluorescent probes using a previously described method with slight modification (Zhuang et al., 2019; Wang et al., 2020). Leaf discs were collected from transgenic and WT tomato plants after salt treatment for 20 d. The leaf discs were soaked in 0.01 mM PBS for 20 min, then placed in 10 μM 2′,7′-dichlorodihydrofluorescein diacetate (Invitrogen, Carlsbad, CA, USA) and incubated under vacuum for 30 min. A confocal laser microscope LSM880 (Carl Zeiss, Oberkochen, Germany) was used to observe the fluorescence signal.

### Transient expression of *MdASG1* in apple fruit

Overexpression vector construction and infiltration of *MdASG1* were conducted as described for the subcellar localization assay. Virus-induced gene silencing was used to silence *MdASG1* in apple fruit. A partial CDS fragment for pTRV2-*MdASG1* (369 bp) was cloned by PCR with specific primers (Supplemental Table S2). *Agrobacterium tumefaciens* containing the target genes was injected into the epidermis of ‘Yinv’ apple fruit with a syringe. *Agrobacterium tumefaciens* carrying the empty vector (PHB or TRV) was used as a control. After infiltration, the fruit were placed in an incubator at 24℃ under a 16 h/8 h (light/dark) photoperiod. After 3 d, the fruit injection sites were sampled for transgene verification and qRT-PCR analysis. After 7 d, the fruit injection sites were sampled for volatile compound analysis using GC-MS. Three biological replicates with at least 15 fruits per group were analyzed.

### Stable overexpression in apple calli and tomato

The full-length CDS of *MdLOX1a* or *MdASG1* was cloned into the PRI 101-AN vector using the primers listed in Supplemental Table S2. The recombinant plasmid was introduced into *A. tumefaciens* strain LBA4404 using the freeze–thaw method. Transformation of apple calli was conducted as described by Zhang et al. (2018). The transgenic calli were used for further analysis. The full-length CDS of *MdASG1* was introduced into the PCB302 vector carrying the CaMV35S promoter and transformed into *A. tumefaciens* strain LBA4404. The primers used for the transformation are listed in Supplemental Table S2. *Agrobacterium* infection solution with OD_600_ = 0.6 was used to infiltrate tomato cotyledons for 15 min. The infiltrated cotyledons were placed on MS medium supplemented with 50 mg·L^−1^ kanamycin to screen for resistant buds. Three lines were confirmed to be transgenic. Tomato fruit harvested at B+7 days from T_3_ transgenic and WT plants were sampled for aroma compound analysis. Three biological replicates with 15 fruits per replicate were analyzed.

### Chlorophyll fluorescence analysis

Chlorophyll fluorescence parameters were measured using a Closed FluorCam FC800 chlorophyll fluorescence imaging system (Photon Systems Instruments, Brno, Czech Republic). Before measurement, the leaves were dark-adapted for 30 min, then analyzed to determine F_0_ (minimum fluorescence) and F_M_ (maximum fluorescence).

### Statistical analysis

Student’s *t*-test (\**P* < 0.05, \*\**P* < 0.01) was used to determine the significance of differences of two samples in this study. Figures were generated using Microsoft Excel. Linear regression analysis was performed using Microsoft Excel and significance of multiple groups was analyzed using Tukey one-way analysis of variance (ANOVA) with SPSS Statistics 22 (IBM Corporation, Armonk, NY, USA).

## Supplemental Data

**Supplemental Figure S1**. Relative expression of *MdLOX* genes during apple fruit development and ripening.

**Supplemental Figure S2**. Linear regression analysis between *MdLOX* expression and the ester content in apple fruit at the ripening stage.

**Supplemental Figure S3**. Fruit of eight apple cultivars harvested at ripening.

**Supplemental Figure S4**. Phylogenetic analysis of plant LOX proteins.

**Supplemental Figure S5**. Background AbA^r^ expression of the yeast Y1H Gold strain containing specific promoters.

**Supplemental Figure S6**. Protein sequence alignment of MdASG1 with AtASG1 and ScASG1.

**Supplemental Figure S7**. Relative expression of stress-related genes in apple calli of the wild-type ‘Orin’ (WT) and *MdASG1*-overexpressing transgenic lines (MdASG1-OE) in response to 50 mM NaCl treatment for 20 d.

**Supplemental Figure S8**. Relative expression of stress-related genes in wild-type (WT) and *MdASG1*-overexpressing (MdASG1-3,6,9) tomato plants in response to 200 mM NaCl treatment for 20 d.

**Supplemental Figure S9**. Transcriptional changes in fatty acid-derived volatile biosynthesis genes in response to 200 mM NaCl treatment in ripening fruit of wild-type (WT) and *MdASG1*-overexpressing (MdASG1) tomato.

**Supplemental Table S1**. Primers used for qRT-PCR analysis in this study.

**Supplemental Table S2**. Primers used to construct or verify vectors in this study.

**Supplemental Table S3**. Probes used for EMSA in this study.

**Supplemental Table S4**. Accession numbers for LOX proteins in the phylogenetic analysis.

## ACKNOWLEDGMENTS

We thank the Shujing Wu Laboratory for providing the plasmids. This work was supported by the National Natural Science Foundation of China (grant no. 31701892 and 31730080) and the Agricultural Variety Improvement Project of Shandong Province (grant no. 2019LZGC007). We thank Liwen Bianji, Edanz Group China (https://www.liwenbianji.cn/ac), for editing the English text of a draft of this manuscript.

